# Clinical and genomic epidemiology of carbapenem-non-susceptible *Citrobacter* spp. at a tertiary healthcare center over two decades

**DOI:** 10.1101/2020.02.21.959742

**Authors:** Ahmed Babiker, Daniel R. Evans, Marissa P. Griffith, Christi L. McElheny, Mohamed Hassan, Lloyd G. Clarke, Roberta T. Mettus, Lee H. Harrison, Yohei Doi, Ryan K. Shields, Daria Van Tyne

**Author notes:** Present address: Department of Pathology and Laboratory Medicine, Emory University School of Medicine, Atlanta, Georgia, USA. Address correspondence to: Daria Van Tyne.

## Abstract

Carbapenem-non-susceptible *Citrobacter* spp. (CNSC) are increasingly recognized as healthcare-associated pathogens. Information regarding their clinical epidemiology, genetic diversity, and mechanisms of carbapenem resistance is lacking. We examined microbiology records of adult patients at the University of Pittsburgh Medical Center (UMPC) Presbyterian Hospital (PUH) from 2000-2018 for CNSC, as defined by ertapenem non-susceptibility. Over this timeframe, the proportion of CNSC increased from 4% to 10% (*P*=0.03), as did carbapenem daily defined doses/1000 patient days (6.52 to 34.5, R^2^=0.831, *P*<0.001), which correlated with the observed increase in CNSC (lag=0 years, R^2^=0.660). Twenty CNSC isolates from 19 patients at PUH and other UPMC hospitals were available for further analysis, including whole-genome short-read sequencing and additional antimicrobial susceptibility testing. Of the 19 patients, nearly all acquired CNSC in the healthcare setting and over half had polymicrobial cultures containing at least one other organism. Among the 20 CNSC isolates, *C. freundii* was the predominant species identified (60%). CNSC genomes were compared with genomes of carbapenem-susceptible *Citrobacter* spp. from UPMC, and with other publicly available CNSC genomes. Isolates encoding carbapenemases (*bla*_KPC-2_, *bla*_KPC-3_, and *bla*_NDM-1_) were also long-read sequenced, and their carbapenemase-encoding plasmid sequences were compared with one another and with publicly available sequences. Phylogenetic analysis of 102 UPMC *Citrobacter* spp. genomes showed that CNSC from our setting did not cluster together. Similarly, a global phylogeny of 64 CNSC genomes showed a diverse population structure. Our findings suggest that both local and global CNSC populations are genetically diverse, and that CNSC harbor carbapenemase-encoding plasmids found in other *Enterobacterales*.

## Introduction

Carbapenem-resistant bacteria have become a major health concern worldwide (1). There are limited therapeutic options for treating infections caused by these multidrug-resistant organisms, resulting in greater morbidity and mortality compared to infections caused by susceptible organisms (2). Furthermore, multidrug-resistant infections place an additional economic burden on healthcare systems (3). In recognition of this threat, the treatment and control of carbapenem-resistant organisms have been prioritized by both the Centers for Diseases Control and Prevention and the World Health Organization (4, 5).

The recent increase in infections caused by carbapenem-resistant organisms in the United States has been largely driven by the dissemination of plasmid-encoded carbapenemases, which are often carried by members of the *Enterobacterales*, particularly *Klebsiella pneumoniae* (6, 7). However, rates of other carbapenem-resistant bacterial species have also increased (8). Among them, carbapenem-non-susceptible *Citrobacter* spp. (CNSC) have become increasingly recognized as a healthcare-associated pathogen (9–13). CNSC isolates have been found to be both genotypically and phenotypically diverse (14, 15), and their resistance to carbapenems is frequently caused by plasmid-encoded carbapenemases, which can be readily acquired through horizontal gene transfer (12, 16).

Information regarding the clinical epidemiology, genetic diversity, and mechanisms of carbapenem resistance among CNSC in the United States are currently limited to a small number of studies and very few isolates (9, 17–19). Here we aimed to investigate the emergence of CNSC within our healthcare system using epidemiology and genomics approaches. We conducted a retrospective analysis of CNSC prevalence and carbapenem use over the last two decades, and compared the genomes of CNSC isolates from our center with other local *Citrobacter* isolates, as well as with CNSC genomes sampled from around the globe. We found that while the CNSC sampled from our center are highly genetically diverse, their diversity is consistent with the local carbapenem-susceptible *Citrobacter* population, as well as with CNSC sampled elsewhere.

## Methods

### Study design and isolate collection

This study was conducted at the University of Pittsburgh Medical Center (UPMC) Presbyterian Hospital (PUH), an adult medical/surgical tertiary care hospital with 762 total beds, 150 critical care unit beds, more than 32,000 yearly inpatient admissions, and over 400 solid organ transplants per year. CNSC isolates were collected from both UPMC-PUH as well as other UPMC hospitals. Ethics approval for this study was obtained from the Institutional Review Board of the University of Pittsburgh.

To investigate CNSC epidemiology at UPMC-PUH, microbiology records of adult patients with a positive clinical culture for CNSC were evaluated from January 1, 2000 to December 31, 2018. Cases were excluded from re-inclusion within 90 days of any CNSC culture. Carbapenem non-susceptibility was defined as non-susceptibility to any carbapenem according to the 2017 Clinical Laboratory Standards Institute (CLSI) interpretative criteria (20). Antibiotic consumption was measured by daily defined doses (DDDs) of any carbapenem (21). To further phenotype and genotype CNSC at UPMC, 20 available CNSC isolates from 19 patients collected between 2013 and 2019 from were included. Isolates were considered community-associated if the organism was isolated from a specimen collected within 72 hours following hospital admission; isolates collected after 72 hours were considered healthcare-associated (22). Clinical characteristics and outcomes of patients with CNSC isolates that underwent further characterization were collected through retrospective chart review. The primary clinical outcome was in-hospital mortality and/or transfer to hospice.

### CNSC isolate characterization

Initial species assignment was performed using standard clinical microbiology laboratory methods, and was confirmed or modified after whole-genome sequencing. Carbapenem non-susceptibility was initially determined by standard clinical microbiology laboratory methods, and was confirmed by the Kirby-Bauer disk diffusion method as per the 2017 Clinical Laboratory Standards Institute (CLSI) interpretative criteria (20). Susceptibility to additional agents was determined by the broth microdilution method (20). Presence of carbapenemase enzyme activity was assessed by modified carbapenem inactivation (mCIM) test (23).

### Genome sequencing and analysis

Genomic DNA was extracted from pure overnight cultures of single bacterial colonies using a Qiagen DNeasy Blood & Tissue Kit according to the manufacturer’s instructions (Qiagen, Germantown, MD). Library construction and sequencing were conducted using the Illumina Nextera NGS Library Prep Kit or the Illumina Nextera XT DNA Library Prep Kit (Illumina, San Diego, CA). Libraries were sequenced on an Illumina NextSeq with 150bp paired-end reads, or an Illumina MiSeq with 300bp paired-end reads. Isolates with suspected plasmid-encoded carbapenemases were sequenced with long-read technology on a MinION device (Oxford Nanopore Technologies, Oxford, United Kingdom). Long-read sequencing libraries were prepared and multiplexed using a rapid multiplex barcoding kit (catalog SQK-RBK004) and sequenced on R9.4.1 flow cells. Base-calling on raw reads was performed using Guppy v2.3.1 (Oxford Nanopore Technologies, Oxford, United Kingdom), and hybrid assembly was performed with both short Illumina reads and long Oxford Nanopore reads using Unicycler v0.4.8beta (24).

Illumina reads were quality filtered and assembled de novo using SPAdes v3.11. Species were identified by Kraken and by performing pairwise comparisons of average nucleotide identity on the assembled genomes using fastANI using the many-to-many method (25). Assemblies were clustered using the hierarchy module of the python package SciPy by single linkage method and a distance criterion of 5% difference in average nucleotide identity. Multi-locus sequence typing (MLST) was performed with the mlst tool at github.com/tseeman/mlst. Genomes were annotated using Prokka v1.13 (26). Core genes were defined using Roary v3.12.0 with a 90% sequence identity cutoff (27). A phylogenetic tree based on a core gene alignment containing 1606 genes identified by Roary was generated using RAxML v8.2.11 (28) by running 1000 bootstrap replicates under the generalized time-reversible model of evolution, a categorical model of rate heterogeneity (GTR-CAT), and Lewis correction for ascertainment bias. The tree was visualized and annotated using Interactive Tree of Life (iTOL) v4 (29). The genomes of closely related isolates were compared with one another with BreSeq (30). AMR gene and plasmid content were assessed by BLASTn of assembled contigs against downloaded ResFinder and PlasmidFinder databases, with 80% sequence identity and 80% sequence coverage cut-off (31). Virulence gene content was assessed using the VirulenceFinder web interface with default settings and the *E. coli* database (32). Carbapenemase-encoding contigs resolved from hybrid assembly of CNSC genomes were annotated using Prokka v1.13 (26), and resistance genes were identified using the ResFinder web interface with default settings (31). CNSC-encoding contigs were compared to one another and to plasmid sequences downloaded from the RefSeq database (n=18,364) using BLASTn of assembled contigs (33, 34). RefSeq and CNSC contigs whose sequences yielded at least 90% coverage of one another in either search direction were aligned to one another using EasyFig (35). Available CNSC genomes were downloaded from the GenBank or Sequence Read Archive repositories maintained by the National Center for Biotechnology Information (NCBI). Genome sequence data generated and analyzed in this study has been deposited in or accessed from SRA/GenBank with accession numbers listed in Supplemental Tables 1, 2, and 3.

### Statistics

The proportion of CNSC was measured by dividing the number of CNSC isolates by the total number of *Citrobacter* spp. isolates tested for carbapenem susceptibility each year. Carbapenem (ertapenem, doripenem, meropenem, and imipenem) DDDs were measured per year at UPMC-PUH (21). Changes in the rate of carbapenem-non-susceptible pathogen isolation over time were measured by linear regression, and comparison with the rate of antibiotic DDDs per year was conducted with time-series cross-correlation analysis. Categorical data were compared using X_2_. Statistical analyses were performed using Stata V15 (StataCorp, College Station, TX) and R V3.5.1 (36).

## Results

### Clinical epidemiology of CNSC

During the study period from 2000 through 2018, 78 unique patients with CNSC were identified from 2817 *Citrobacter* spp. isolates tested. *Citrobacter* spp. were the seventh most common carbapenem-non-susceptible gram-negative bacteria, and fifth most common carbapenem-non-susceptible *Enterobacterales* at our center during this time period. The proportion of *Citrobacter* spp. isolates that were CNSC increased significantly over time (R^2^=0.257, *P*=0.03), from 4% in 2000 to 10% in 2018 (Figure 1). Daily defined doses (DDDs) of carbapenems per 1000 patient days also increased during the same time period, from 6.52 in 2000 to 34.5 in 2018 (R^2^=0.831, *P*<.001). We found that the increase in DDDs correlated with the increase in CNSC over the same time period (lag= 0 years, R^2^=0.660) (Figure 1).

**Figure 1.**
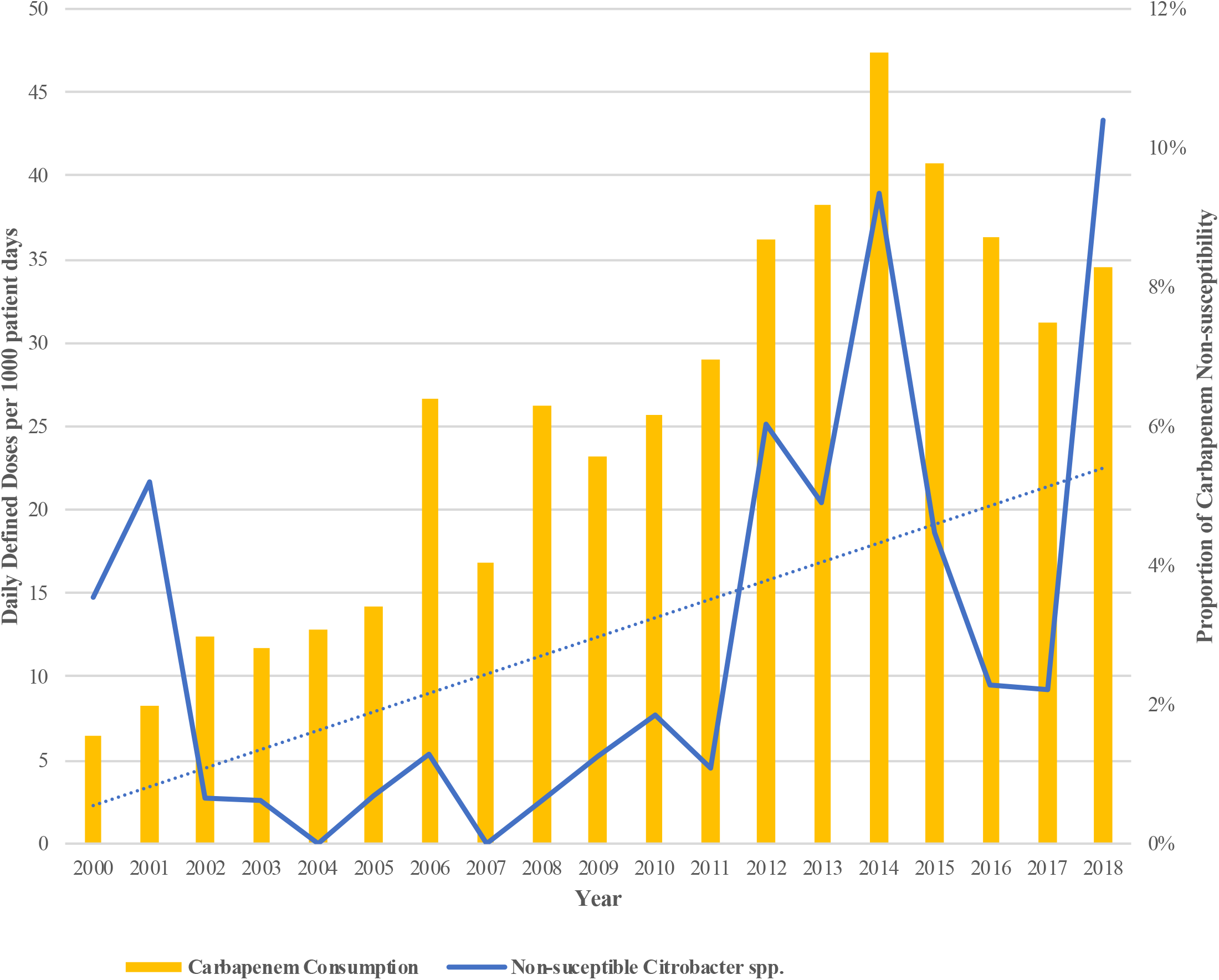
Carbapenem consumption and proportion of carbapenem-non-susceptible *Citrobacter* spp. (CNSC), 2000-2018. Carbapenem daily defined doses (DDDs) per 1000 patient days (yellow bars) and the proportion of *Citrobacter* spp. isolates that were carbapenem-non-susceptible (solid blue line) were quantified for each year between 2000 and 2018 at UPMC PUH. Of 2817 total *Citrobacter* spp. isolates tested, 78 unique patients had CNSC (defined as ertapenem non-susceptible). The dotted blue line shows a linear regression for increased CNSC proportion over time (R^2^=0.257, P=0.03). Carbapenem DDDs per 1000 patient days also increased over time (R^2^=0.831, P<.001), and correlated with the increase in CNSC (lag=0 years, R^2^=0.660).

### Isolation and characterization of CNSC

Twenty CNSC isolates from 19 patients from UPMC-PUH and two additional UPMC hospitals were available for further analysis (Table 1). Among these patients, the median age was 65 (range, 26-92) and 37% were female (7/19). The majority of patients had multiple comorbidities, frequently acquired CNSC in the healthcare setting (84%, 16/19), had polymicrobial cultures (57%, 11/19), and had high rates of in-hospital mortality/discharge to hospice (47%, 9/19) (Table 1). We sequenced the genomes of all 20 CNSC isolates on the Illumina platform (Supp. Table 1), and constructed a phylogenetic tree that also included an additional 82 carbapenem-susceptible *Citrobacter* spp. isolates collected by the Enhanced Detection System for Hospital-Associated Transmission (EDS-HAT) project (Figure 2) (37, 38) (Supp. Table 4). Among the 20 CNSC isolates, *C. freundii* was the predominant species (60%, 12/20), followed by *C. werkmanii* (20%, 4/20), *C. koseri* (10%, 2/20), and *C. farmeri* (5%, 1/20). One CNSC isolate, YDC693, was originally identified as *C. freundii* but only showed 90-92% average nucleotide identity to other *C. freundii* genomes (Supp. Table 4). This isolate appears to belong to a new, unnamed *Citrobacter* species. YDC693 (*Citrobacter* sp.) and YDC697-2 (*C. farmeri*) were both cultured from the same patient, and were sampled approximately two weeks apart. Their distribution throughout the genome phylogeny suggested that the CNSC isolates were largely genetically distinct from one another (Figure 2, Supp. Table 4). The one exception was RS259 and YDC849-1, which were found to have fewer than 20 genetic variants (single nucleotide polymorphisms and insertion/deletion variants) that distinguished them from one another, despite being isolated from patients at two different facilities (Facility A vs. Facility B).

**Table 1:** Clinical characteristics of patients with CNSC

| Isolate ID | Age | Gender | Year | Source | Facility | CNSC Organism | Culture site | Clinical Syndrome | Polymicrobial culture (Y/N) | Other organisms | Comorbid Conditions | Outcome of hospitalization |
| --- | --- | --- | --- | --- | --- | --- | --- | --- | --- | --- | --- | --- |
| RS77 | 49 | M | 2018 | HA | A | *C. freundii* | Rectal swab | Colonization | N | None | ESLD, liver transplant | Discharge to home |
| RS102 | 74 | F | 2018 | HA | A | *C. werkmanii* | Rectal swab | Colonization | N | None | ESLD, liver transplant | In-hospital death |
| RS189 | 66 | F | 2017 | HA | A | *C. werkmanii* | BAL | Pneumonia | N | None | Diverticulosis | Discharge to facility |
| RS226 | 56 | F | 2018 | HA | A | *C. freundii* | Urine | Colonization | Y | *E. coli* | Opiate dependency, CHF, HTN | In-hospital death |
| RS236 | 70 | M | 2018 | HA | A | *C. freundii* | Peritoneal fluid | Intra-Abdominal | Y | *C. freundii* (ESBL), *E. faecium* (VRE), *C. tropicalis*, *C. parapsilosis* | Pancreatic cancer, COPD, DM, HTN | Transfer to hospice |
| RS237 | 80 | F | 2018 | Community | A | *C. freundii* | Urine | Colonization | N | None | Sinus OM | Discharge to home |
| RS259 | 61 | F | 2018 | HA | B | *C. freundii* | Peritoneal fluid | Intra-Abdominal | Y | *E. coli*, *P. aeruginosa* | Metastatic lung cancer, COPD, DM | In-hospital death |
| RS289 | 92 | M | 2018 | Community | A | *C. koseri* | Urine | UTI | Y | *E. faecalis* | Bladder cancer, dementia, CKD, COPD, CHF | Discharge to home |
| YDC608 | 57 | M | 2013 | HA | A | *C. freundii* | BAL | Colonization | N | None | HLD | Transfer to hospice |
| YDC638-2 | 27 | M | 2013 | HA | A | *C. freundii* | Biliary drainage | Intra-Abdominal | Y | *E. faecium* (VRE) | ESLD, Crohn's diseases, PSC, liver transplant | Discharge to home |
| YDC645 | 67 | F | 2013 | HA | C | *C. freundii* | Blood | SSTI/Endocarditis | Y | *Bacteroides* sp., *E. raffinosus* | CAD, CHF, DM, ESRD, dementia | Transfer to hospice |
| YDC661 | 64 | M | 2014 | HA | A | *C. freundii* | BAL | Pneumonia | Y | *S. maltophilia* | Heart transplant | Discharge to facility |
| YDC667-1 | 73 | M | 2014 | HA | A | *C. werkmanii* | BAL | Pneumonia | Y | *K. pneumoniae* (ESBL) | CHF, DM, CAD, CKD | In-hospital death |
| YDC689-2 | 61 | M | 2015 | HA | A | *C. koseri* | BAL | Pneumonia | N | None | ESLD, liver transplant | Discharge to facility |
| YDC693* | 65 | M | 2015 | HA | A | *Citrobacter* spp. | BAL | Pneumonia | Y | *E. coli* (ESBL) | SBT, adrenal insufficiency | In-hospital death |
| **YDC697-2\*** | 65 | M | 2015 | HA | A | *C. farmeri* | Tracheostomy site drainage | SSTI | Y | *K. oxytoca* (KPC) | SBT, adrenal insufficiency | In-hospital death |
| **YDC730** | 71 | M | 2015 | HA | D | *C. werkmanii* | Pelvic abscess | Intra-Abdominal | Y | *E. faecium* (VRE) | Multiple myeloma | Discharge home |
| **YDC849-1** | 26 | F | 2018 | HA | A | *C. freundii* | Urine | UTI | Y | *C. freundii* (NDM) | CVID | In-hospital death |
| **YDC876** | 53 | M | 2019 | HA | A | *C. freundii* | BAL | Pneumonia | N | None | CAD, CHF | Discharge home |
| **YDC880** | 73 | M | 2019 | HA | A | *C. freundii* | Rectal swab | Colonization | N | None | Lung transplant | In-hospital death |

Abbreviations: BAL: Bronchoalveolar lavage, CAD: coronary artery disease, CHF: congestive heart failure, CKD: chronic kidney disease, COPD: chronic pulmonary disease, CNSC: carbapenem non-susceptible *Citrobacter spp*, CVID: common variable immunodeficiency, ESBL: Extend-spectrum beta-lactamase, ESLD: end-stage liver disease, ESRD: end-stage renal disease DM: diabetes mellitus, F: female, HAP: hospital-acquired pneumonia, HCC: hepatocellular carcinoma, HTN: hypertension, KPC: *Klebsiella pnuemoniae* carbapenemase, OM: osteomyelitis, M: male, NDM: New Delhi metallo-beta-lactamase, PSC: primary sclerosing cholangitis, SBT: small bowel transplant, SSTI: skin and soft tissue infection, VRE: Vancomycin-resistant *Enterococcus*, UTI: urinary tract infection
\*Same patient isolates

**Figure 2.**
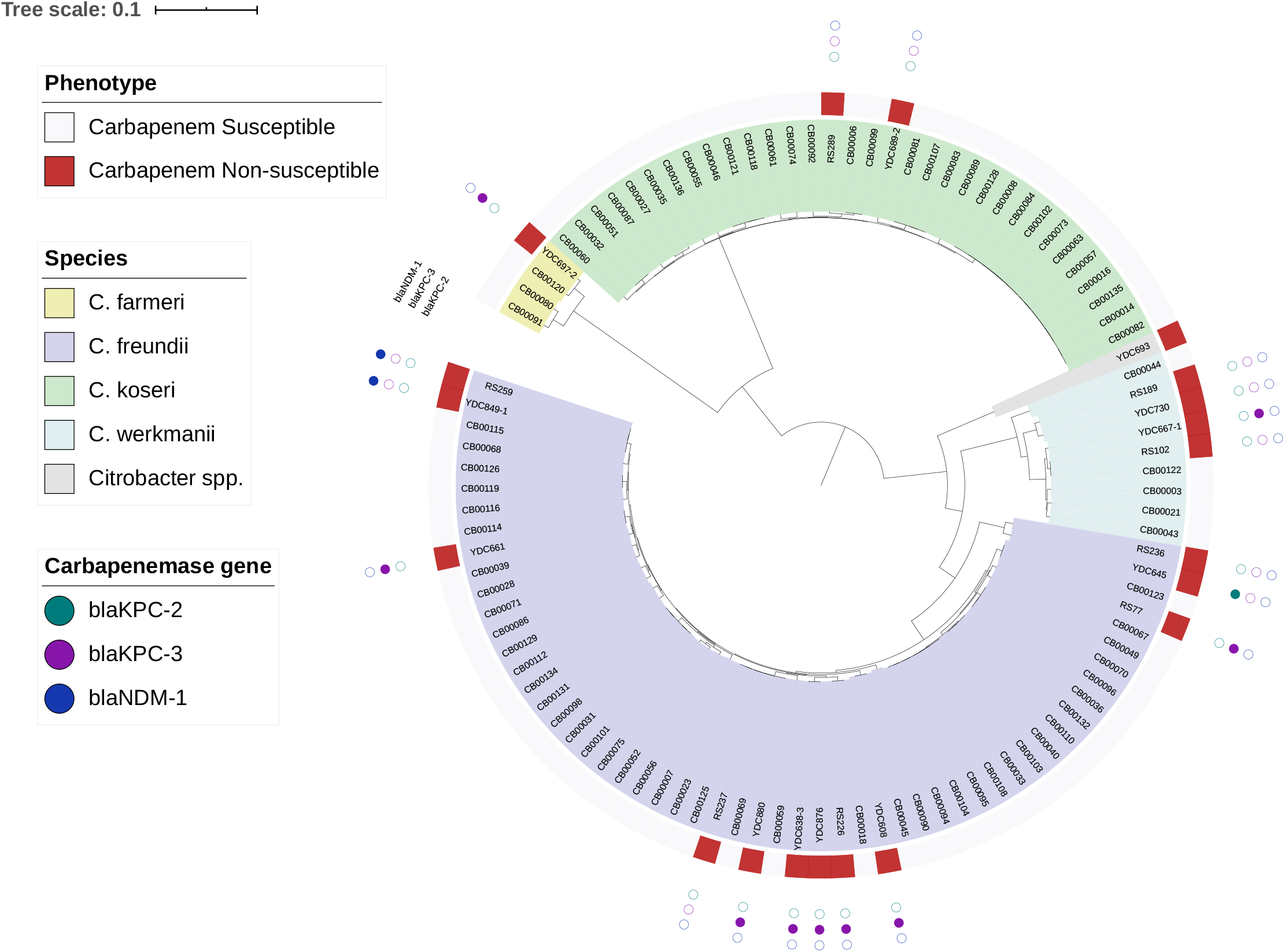
Local phylogeny of carbapenem-susceptible and non-susceptible *Citrobacter* spp. from UPMC. A phylogenetic tree of 102 local *Citrobacter* spp. genomes (82 carbapenem-susceptible and 20 carbapenem-non-susceptible isolates) was generated based on an alignment of 1606 core genes using RAxML (20). The tree was visualized and annotated using Interactive Tree of Life (iTOL) (28). The tree is annotated based on carbapenem susceptibility phenotype, species, and carbapenemase genes identified, if any, in the genome of each isolate.

### Antimicrobial susceptibility and identification of carbapenemase and virulence genes

While the CNSC isolates we collected were originally defined as non-susceptible to ertapenem, they displayed variable susceptibility patterns to other carbapenem antibiotics. Only about half of the isolates (55%, 11/20) were non-susceptible to meropenem, with three isolates having intermediate resistance and eight being resistant (Table 2). We tested all isolates for the presence of a carbapenemase using a modified carbapenem inactivation (mCIM) test (23), for which 13 isolates (65%) tested positive. Carbapenemase genes were present in the genomes of all 13 isolates (Table 2). Among the carbapenemases identified, *bla*_KPC-3_ was predominant (10/13), followed by *bla*_NDM-1_ (2/13) and *bla*_KPC-2_ (1/13). We also tested all 20 CNSC isolates against novel β-lactam/β-lactamase inhibitor agents, and found they were frequently susceptible to ceftazidime-avibactam (17/20, 85%) and meropenem-vaborbactam (18/20, 90%). As expected, both isolates with *bla*_NDM-1_ exhibited phenotypic resistance to both agents (Table 2). Isolate RS237 was also found to be resistant to ceftazidime-avibactam, even though it did not encode a carbapenemase enzyme. We compared RS237 with the most closely related carbapenem-susceptible isolate, CB00023, and found a large number of mutations separating them from one another (Supp. Table 5). One of these was a missense mutation (S219I) in *acrE*, which is predicted to encode a multidrug export protein and could be a candidate resistance-associated gene.

**Table 2:** Carbapenemase genes and antimicrobial susceptibilities to novel beta-lactam/beta-lactamase inhibitor agents

| Isolate | Organism | Carbapenemase gene | Meropenem MIC | Ceftazidime-Avibactam MIC | Meropenem-Vaborbactam MIC | Modified Carbapenemase Inactivation Test Result |
| --- | --- | --- | --- | --- | --- | --- |
| RS77 | *C. freundii* | *blaKPC-3* | 1 | 0.5 | 0.015 | Positive |
| RS102 | *C. werkmanii* | - | 0.25 | 0.5 | 0.06 | Negative |
| RS189 | *C. werkmanii* | - | ≤0.06 | 0.5 | 0.015 | Negative |
| RS226 | *C. freundii* | *blaKPC-3* | 2 | <0.25 | 0.015 | Positive |
| RS236 | *C. freundii* | - | 2 | 2 | 0.5 | Negative |
| RS237 | *C. freundii* | - | 4 | 64 (R) | 0.5 | Negative |
| RS259 | *C. freundii* | *blaNDM-1* | 16 | >256 (R) | >8 (R) | Positive |
| RS289 | *C. koseri* | - | 0.12 | 2 | 0.12 | Negative |
| YDC608 | *C. freundii* | *blaKPC-3* | 16 | 4 | 0.06 | Positive |
| YDC638-3 | *C. freundii* | *blaKPC-3* | 2 | 2 | 0.06 | Positive |
| YDC645 | *C. freundii* | *blaKPC-2* | ≤0.06 | <0.25 | 0.03 | Positive |
| YDC661 | *C. freundii* | *blaKPC-3* | 1 | 4 | 0.06 | Positive |
| YDC667-1 | *C. werkmanii* | *blaKPC-3* | 1 | 0.5 | 0.03 | Positive |
| YDC689-2 | *C. koseri* | - | 0.5 | 4 | 0.12 | Negative |
| YDC693* | *Citrobacter* sp. | *blaKPC-3* | 16 | 1 | 0.03 | Positive |
| YDC697-2* | *C. farmeri* | *blaKPC-3* | 32 | 4 | 0.12 | Positive |
| YDC730 | *C. werkmanii* | - | 0.12 | 0.5 | 0.12 | Negative |
| YDC849-1 | *C. freundii* | *blaNDM-1* | 16 | >256 (R) | 16 (R) | Positive |
| YDC876 | *C. freundii* | *blaKPC-3* | 8 | 1 | 0.06 | Positive |
| YDC880 | *C. freundii* | *blaKPC-3* | 4 | 1 | 0.03 | Positive |

All isolates were determined to be Carbapenem non-susceptible based on ertapenem non-susceptibility.
MICs are reported as ug/mL and resistance (R) designations are based on CLSI breakpoints.
\* Same patient isolates.

In addition to carbapenemase genes, we also compared the acquired antibiotic resistance gene content between CNSC and carbapenem-susceptible EDS-HAT isolates (Supp. Table 6). CNSC isolate genomes often carried genes encoding resistance to aminoglycoside, β-lactam, fluoroquinolone and tetracycline antibiotic classes. The average number of resistance genes was significantly higher among the CNSC isolates compared to carbapenem-susceptible EDS-HAT isolates (mean (sd), 8 (5) vs. 3 (20); *P* <0.001); Supp. Table 6). Finally, we identified three CNSC isolates with virulence genes previously described in *E. coli* (32). RS289 and YDC689-2 both encoded a *senB* gene (Genbank accession: CP000038), which encodes an enterotoxin (39). Additionally, YDC667-1 encoded an *astA* gene (Genbank accession: AF411067) which encodes an EAST-1 heat-stable toxin (40). These genes were also found among carbapenem-susceptible isolates (Supp. Table 7).

### Global phylogeny of CNSC

To understand how the genomic diversity of the study isolates compared to CNSC isolates from other locations, we searched the NCBI databases for additional publicly available CNSC genomes. Using search terms “Citrobacter” and “Carbapenem,” we identified 64 additional CNSC genomes (Supp. Table 3). A global phylogeny of these 64 CNSC genomes combined with the 20 from this study showed abundant genetic heterogeneity (Figure 3). We investigated the species distribution of the global CNSC population using fastANI, and found that, similar to our UPMC isolates, the global CNSC population was dominated by *C. freundii* (41/64, 64%), followed by *C. amalonaticus* (9/64, 14%), *C. werkmanii* (3/64, 5%), and *C. koseri* (2/64, 3%). Carbapenem-non-susceptible *C. amalonaticus* was not found among our UPMC isolates, but has been isolated from the United States, South America, and Europe (Figure 3, Supp. Table 3). *Citrobacter* sp. YDC693 was found to cluster with an additional three isolates from the United States (Supp. Table 3). Three other global CNSC (two from the United States and one from China) appeared to belong to another distinct *Citrobacter* species with 90-93% average nucleotide identity to *C. freundii*. The proportion of global CNSC isolate genomes that encoded carbapenemase enzymes (49/64, 77%) was similar to our UPMC isolate set (Figure 3), however the diversity of enzyme types was greater and included *bla*_NDM-5_, *bla*_IMP-38_, and *bla*_OXA-48_-like enzymes.

**Figure 3.**
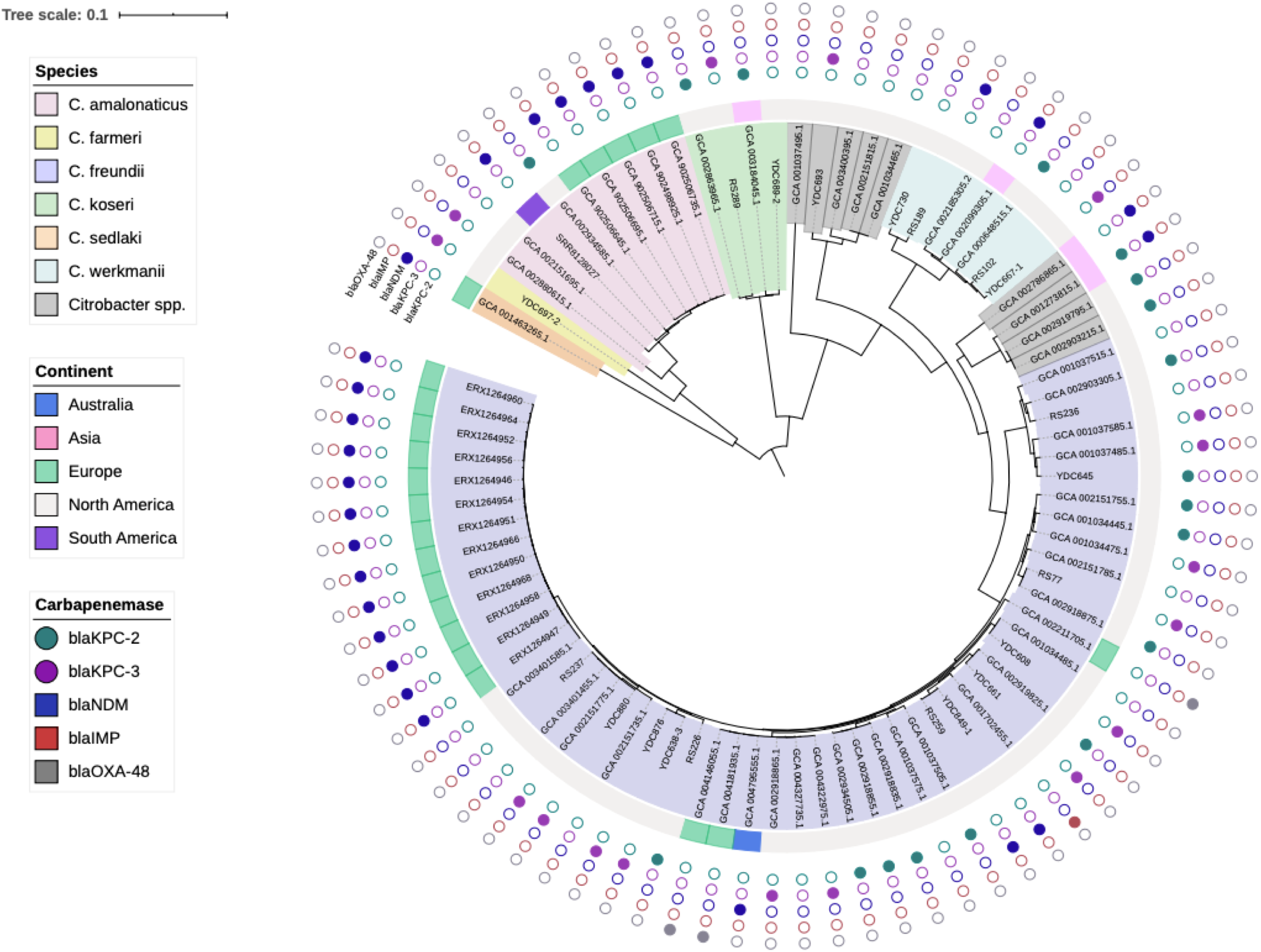
Global phylogeny of available CNSC genomes. A phylogenetic tree of 84 CNSC genomes (20 from this study and 64 from the NCBI) was generated based on an alignment of 1842 core genes using RAxML (20). The tree was visualized and annotated using Interactive Tree of Life (iTOL) (28). The tree is annotated based on species, continent of isolation, and carbapenemase genes identified, if any, in the genome of each isolate.

### Carbapenemase-encoding plasmid diversity

To better understand the genetic context of the carbapenemase enzymes encoded by our UPMC CNSC isolates, we conducted Oxford Nanopore long-read sequencing and hybrid assembly of the 13 carbapenemase-carrying isolate genomes (Table 3). Complete, circular plasmids were resolved for seven isolates, and six of these contained replicons belonging to the *IncA/C2*, *IncL/M*, *IncN*, and unnamed *repA* families. The assembly of RS259 contained a 21.4 kb circular contig that encoded the *bla*_NDM-1_ carbapenemase but lacked a readily identifiable plasmid replicon. This circular contig was highly similar to a region of the 161 kb, *bla*_NDM-1_-encoding plasmid resolved from the YDC849-1 genome, suggesting that the *bla*_NDM-1_-encoding integron may have excised from the surrounding plasmid sequence in RS259. Additionally, the YDC876 genome contained two plasmids of different sizes with distinct replicons that both harbored *bla*_KPC-3_ carbapenemase genes. Direct comparison of these two plasmids confirmed they were distinct, even though they encoded the same carbapenemase and other acquired antimicrobial resistance genes (Table 3). Finally, five carbapenemase-encoding contigs were not completely resolved by hybrid assembly; the YDC638-3_4 contig was more than 187 kb long and contained an IncA/C2 replicon along with *bla*_KPC-3_, and we suspect that it constitutes a nearly complete plasmid sequence. In the other four cases, carbapenemase-encoding contigs were short (less than 20kb) and the plasmids in these isolates could potentially be resolved with additional long-read sequencing.

**Table 3:** Carbapenemase-encoding contigs identified in CNSC genomes

| Isolate_<br>Contig | Length<br>(bp) | Circular<sup>1</sup> | Replicons | Carbapenemase<br>gene | Additional Acquired Antimicrobial Resistance Genes<sup>2</sup> |
| --- | --- | --- | --- | --- | --- |
| RS77_21 | 10,011 | No | None | *bla<sub>KPC-3</sub>* | None |
| RS226_33 | 4,828 | No | None | *bla<sub>KPC-3</sub>* | None |
| RS259_9 | 21,420 | Yes | None | *bla<sub>NDM-1</sub>* | *aac(3)-IId, aac(6')-Ib-cr, aadA16, ARR-3, catB3, dfrA27, mph(A), sul1* |
| YDC608_5 | 172,511 | Yes | IncA/C2 | *bla<sub>KPC-3</sub>* | *aac(6')-Ib, aadA1, ant(2'')-Ia, bla<sub>OXA-9</sub>, bla<sub>SHV-7</sub>, bla<sub>TEM-1A</sub>* |
| YDC638-3_4 | 187,424 | No | IncA/C2 | *bla<sub>KPC-3</sub>* | *aac(6')-Ib, aadA1, ant(2'')-Ia, bla<sub>OXA-9</sub>, bla<sub>TEM-1A</sub>, qnrA1* |
| YDC645_3 | 44,364 | Yes | repA | *bla<sub>KPC-2</sub>* | None |
| YDC661_9 | 15,031 | No | None | *bla<sub>KPC-3</sub>* | None |
| YDC667-1_41 | 4,894 | No | None | *bla<sub>KPC-3</sub>* | None |
| YDC693_4* | 258,721 | Yes | IncA/C2,<br>IncN | *bla<sub>KPC-3</sub>* | *aac(6')-Ib, aadA1, bla<sub>OXA-9</sub>, bla<sub>SHV-7</sub>, bla<sub>TEM-1A</sub>, dfrA14* |
| YDC697-2_6* | 62,530 | Yes | IncN | *bla<sub>KPC-3</sub>* | *dfrA14* |
| YDC849-1_2 | 160,983 | Yes | IncA/C2 | *bla<sub>NDM-1</sub>* | *aac(3)-IId, aac(6')-Ib-cr, aadA16, aph(3'')-Ib, aph(3')-Ia, aph(6)-Id, ARR-3, catB3, dfrA27, floR, mph(A), qnrS1, sul1, sul2, tet(A)* |
| YDC876_2 | 176,497 | Yes | IncA/C2 | *bla<sub>KPC-3</sub>* | *aac(6')-Ib, aac(6')-Ib-cr, aadA1, ant(2'')-Ia, bla<sub>OXA-9</sub>, bla<sub>SHV-7</sub>, bla<sub>TEM-1A</sub>, qnrA1, sul1* |
| YDC876_4 | 78,220 | Yes | IncL/M | *bla<sub>KPC-3</sub>* | *aac(6')-Ib, aac(6')-Ib-cr, aadA1, bla<sub>OXA-9</sub>, bla<sub>TEM-1A</sub>* |
| YDC880_4 | 88,095 | Yes | IncL/M | *bla<sub>KPC-3</sub>* | *aadA2, bla<sub>SHV-30</sub>, dfrA12, sul1* |

<sup>1</sup>Circular contigs were identified through hybrid assembly with Unicycler (24).
<sup>2</sup>Antimicrobial resistance genes were identified by querying the ResFinder database (29).
\* Contigs from same patient isolates.

To determine how the carbapenemase-encoding plasmids in this study compared to one another and whether they were unique to our study, we conducted pairwise comparisons of the resolved plasmids. In addition, we searched the RefSeq database (34) for plasmids that showed substantial homology and high sequence identity to one or more of the CNSC plasmids from our UPMC isolates (Figure 4). Three of the plasmids we identified (YDC608_5, YDC876_2, and YDC638-3) were highly similar to one another, and were found in *C. freundii* isolates belonging to two different sequence types (ST185 and ST116, Figure 4A). While YDC608 and YDC876 belonged to two different *C. freundii* sequence types, their plasmids were more similar to one another than the plasmid from YDC638-3, which belonged to the same sequence type as YDC876. These plasmids were also highly similar to the pCAV1193-166 plasmid found in a *bla*_KPC_-carrying *K. pneumoniae* isolate from Virginia (Figure 4A) (41). Separately, the YDC849-1_2 plasmid encoding *bla*_NDM-1_ had high similarity with plasmid p1540-2, which was found in a carbapenem-resistant *E. coli* isolate from Hong Kong (GenBank accession: CP019053.1) (Figure 4B). Finally, the *bla*_KPC-3_-carrying plasmids YDC693_4 and YDC697-2_6 were from isolates of different species that came from the same patient. Despite being different sizes (259kb vs. 63kb), the plasmids showed some similarity to one another (Figure 4C). These data suggest possible transfer of a *bla*_KPC-3_-encoding mobile element between the isolates from this patient, however independent acquisition or independent transfer from another species cannot be ruled out.

**Figure 4.**
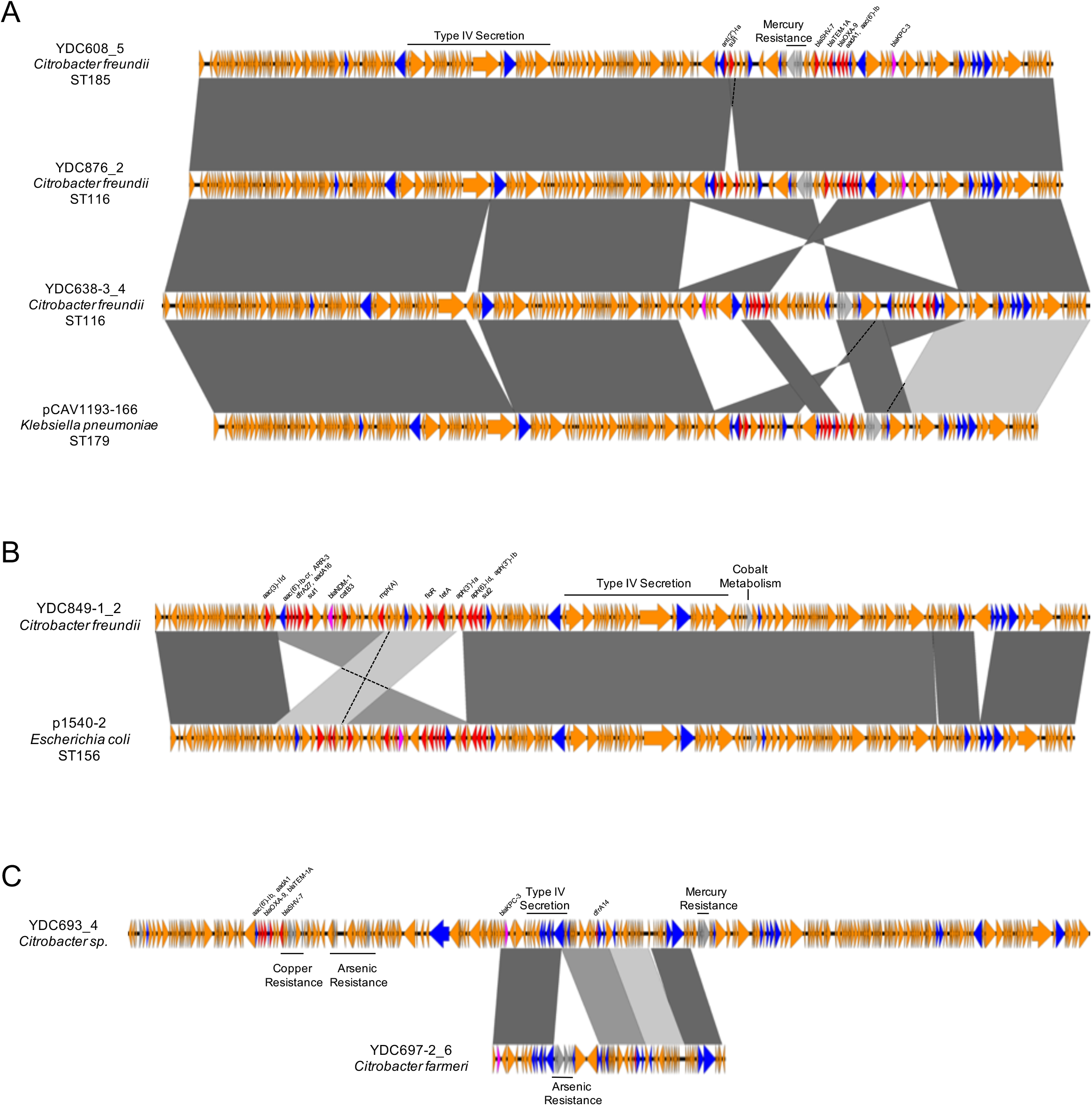
Carbapenemase-encoding plasmid diversity among and between CNSC genomes. Genome contigs encoding carbapenemases and plasmid replicons were compared to each other, and with sequences deposited in the National Center for Biotechnology Information (NCBI). Sequences were aligned to one another with EasyFig. Sequence names correspond to isolateID_contig# from hybrid assembly, or to the sequence name from NCBI. Species and sequence type (ST) are listed, where available. ORFs are colored by function (blue = mobilization, pink = carbapenemase, red = other antibiotic resistance, gray = metal-interacting, orange = other/hypothetical). Antibiotic resistance genes, metal-interacting operons, and Type IV secretion system components are labeled. Grey blocks between sequences indicates regions >5kb with >98% nucleotide identity, with darker shading indicating higher identity.

## Discussion

In this study, we conducted a retrospective review of the clinical and genomic epidemiology of CNSC over the past two decades at a large healthcare center in the United States. We analyzed the genomes of 20 CNSC and 82 carbapenem-susceptible *Citrobacter* spp. sampled locally, as well as 64 publicly available genomes sampled from around the globe. We found that the rates of CSNC increased significantly over the last two decades at our center, that CSNC were frequently acquired in the healthcare setting along with other healthcare-associated organisms, and that patients from whom CNSC were isolated often had poor clinical outcomes. Our phylogenetic analyses revealed genetically diverse CNSC populations both locally and globally, suggesting that CNSC most often arise independently from one another. We also found that carbapenem-non-susceptibility was often mediated by acquisition of carbapenemases genes, with *bla*_KPC-3_ being the predominant carbapenemase identified among CNSC isolates in our setting.

*Citrobacter* spp. have become increasing recognized as a cause of multidrug-resistant healthcare-associated infections around the world (10, 42–44), and prior reports have identified CNSC predominantly from healthcare sources and often associated with nosocomial outbreaks (9, 12, 45). We detected a significant increase in the proportion of *Citrobacter* spp. isolates that were carbapenem-non-susceptible over the last two decades, which correlated with increased use of carbapenems at our center. While the incidence of carbapenem-resistant organisms has increased worldwide over recent years (7, 46), attention has been largely focused on other carbapenem-resistant members of the *Enterobacterales*, such as *Enterobacter* spp., *E. coli*, and *Klebsiella* spp. (7). As with other gram-negative species, increasing antibiotic resistance among *Citrobacter* spp. is of significant concern, as our findings show that many CNSC appear to have acquired resistance genes from other bacteria by horizontal gene transfer.

Our analysis revealed extensive genomic diversity among both CNSC and carbapenem-susceptible *Citrobacter* sampled from our center. This is similar to previous analyses of CNSC from non-US centers that used more classical molecular typing methods (14, 47, 48). Among *Citrobacter* species*, C. freundii* is most commonly associated with both clinical disease (44) and multidrug-resistant phenotypes (14, 15, 49). Our findings were consistent with these prior reports – *C. freundii* was the most frequent CNSC species we observed, though we also found CNSC belonging to four additional species. The global CNSC population was similarly diverse, but *C. freundii* was again the most prevalent species observed, which may be due to its higher rate of antibiotic resistance compared to other *Citrobacter* spp. (43, 44).

The term “carbapenem non-susceptibility” encompasses a wide range of phenotypic susceptibilities, which can be caused by different mechanisms. Among the CNSC isolates we collected, roughly two thirds were found to produce carbapenemases, a rate that is similar to prior reports (14), and to the global CNSC population. CSNC have been found to encode a diverse array of carbapenemases. For example, a study by Arana *et al*. found five different carbapenemase types among *Citrobacter* spp. isolates collected from Spain (14), and similar results were also demonstrated in a study of carbapenemase-producing *Enterobacterales* in China (15). It has been suggested that carbapenemase diversity depends on local geography (46), and future studies of larger populations may confirm or refute this notion. The high diversity of carbapenemase-encoding plasmids we found, all from isolates of a single genus at a single hospital, highlights the complexity of antibiotic resistance gene transfer between pathogens in the hospital setting (50). Even with a relatively small number of isolates, we observed closely related plasmids in genetically distinct bacteria, identical *bla*_KPC-3_-encoding mobile elements on different plasmids carried by the same bacterial isolate, and similar carbapenemase-encoding plasmids in CNSC of different species that were isolated from the same patient. These findings underscore the highly dynamic and variable transfer of carbapenemase plasmids into and among CNSC isolates, and suggest that carbapenemase-encoding plasmids may not be stably maintained in the absence of strong selective pressure.

There were several limitations to this study. While our study presents the largest genomic analysis of CNSC from the United States, the number of isolates we included was still rather limited. Furthermore, the correlation between carbapenem consumption and proportion of CNSC is a strictly ecologic analysis. Additionally, our genomic analysis of resistance determinants was limited to acquired carbapenemase genes, and we did not investigate additional resistance mechanisms such as chromosomal *ampC* genes, efflux pumps, or outer membrane protein mutations, which are known to be associated with carbapenem resistance. Furthermore, many of the global isolate genomes we analyzed were from the United States and/or were part of outbreak investigations, thus they may not be representative of the true global diversity of CNSC. Additionally, we only made *in silico*, sequence-based comparisons of the plasmids we resolved; as such, we cannot comment on their capacity for conjugative transfer. Finally, we were unable to determine whether the poor clinical outcomes among the patients from whom CNSC were isolated were indeed attributable to CNSC infection.

As they become more prevalent in the healthcare system, further studies will be needed to increase our understanding of CNSC genomic diversity and resistance mechanisms. In particular, examining the local and global epidemiology of horizontal transfer of drug resistance elements among *Citrobacter* spp. – and between *Citrobacter* and other *Enterobacterales* species – would provide valuable insights into risk factors and other trends that could be targeted to limit the occurrence and spread of CNSC.

## Acknowledgements

We gratefully acknowledge Hayley Nordstrom, Daniel Snyder, and Vaughn Cooper for generating genome sequence data. This study was funded by the National Institute of Allergy and Infectious Diseases (R21AI109459 and R01AI127472 to L.H.H.), and by the University of Pittsburgh Department of Medicine. The funders had no role in study design, data collection and interpretation, or the decision to submit the work for publication.

**Preliminary data included in this work was presented at the ID Week 2019 conference (abstract #485).**

